# Research on Mitochondrial DNA Mutations in Patients with SCA3/MJD

**DOI:** 10.1101/101238

**Authors:** Zhen Liu, Jie Zhou, Xiaomeng Yin, Shuying Shi, Weining Sun, Hong Jiang, Lu Shen, Beisha Tang, Junling Wang

## Abstract

Spinocerebellar ataxia type 3 (SCA3) is a degenerative neurological disorders caused by trinucleotide repeat expansion within the ataxin-3 gene. It is characterized by multi-system involvement and diverse clinical phenotypes, which cannot be fully explained the length of the CAG repeats. One possible explanation for the phenotypic heterogeneity could be the presence of mitochondrial DNA mutations that modify disease severity. To explore the role of Mitochondrial DNA(mtDNA) variations in SCA3 pathogenesis, we analyzed polymorphisms of six mitochondrial genes, *MT-LT1, MT-ND1, MT-CO2, MT-TK, MT-ATP8* and *MT-ATP6*, in 102 unrelated SCA3/MJD patients and 100 healthy controls. The results showed that there were 24 variations of those mtDNA genes in the SCA3 patients and only 10 in the unrelated healthy controls. There was no difference of the relative mtDNA copy number variation between the SCA3 patients and healthy controls (93.20 vs. 89.66, P>0.05). In the group of SCA3 patients, the relative mtDNA copy number showed a negative correlation between the number of CAG repeats (r=−0.210, P < 0.05), but did not correlate with the age at diagnosis, the age of onset, disease duration, ICARS scores and SARA scores. Our research demonstrated that the frequency of mutated mtDNA in SCA3 patients was higher than that in the healthy group. The mtDNA relative copy number in SCA3 patients was not significantly different compared to the healthy group. Thus, the copy number might not be treated as a biomedical indicator when measuring the severity of illness in SCA3 patients.

## Introduction

Spinocerebellar ataxia type 3 (SCA3), also known as Machado-Joseph Disease (MJD), is the most common form of SCA worldwide and characterized by progressive cerebellar ataxia of the gait and limbs, dysarthria, dysphagia and pyramidal signs. Meanwhile, SCA3/MJD is variably associated with other symptoms such as nystagmus, muscle atrophy, dystonia, extrapyramidal signs and autonomic dysfunction. The expansion of a CAG tract in the coding region of the SCA3-causative gene *ATXN3* (chromosome 14q32.1) translates into an expanded polyglutamine tract at the C-terminus of the ataxin-3 protein. There is consensus that the toxic gain-of-function mutant ataxin-3 plays a fundamental role in SCA3 pathogenesis(Yu *et al.* 2009; Nobrega *et al.* 2013). However, the distinct pathogenic mechanisms of the CAG repeat expansion remain unclear, and toxicity of the widely expressed polyglutamine protein cannot fully explain the selective neuronal degeneration seen in this disease(Maruyama *et al.* 1995; Padiath *et al.* 2005; Yu *et al.* 2009; Ramos *et al.* 2015).

Mitochondrial DNA, which exists completely apart from the nuclear genome, contains 16,569 base pairs and codes for 37 genes. These genes are all necessary for the synthesis of mitochondrial proteins. Mitochondrial dysfunction can decrease ATP production, increase toxic free radicals and disrupt cytoplasmic calcium balance, leading to excitotoxicity and apoptosis(Petrozzl *et al.* 2007). Mitochondrial dysfunction associated with several degenerative neurological diseases(Waldbaum and Patel 2010; Pareyson *et al.* 2014).

Nuclear DNA or mtDNA mutations can both cause mitochondrial dysfunction. Several groups have explored the relationship between spinocerebellar ataxia and mutations in mitochondrial DNA, such as Casali(Casali *et al.* 1999), Chinnery(Chinnery *et al.* 2002) and Bing-Wen Soong(Lee *et al.* 2007) and have found mtDNA single nucleotide variants (SNVs) and relative mtDNA copy number variations in SCA families and some sporadic patients. Based on these findings, we wondered whether mtDNA mutations might play a role in SCA3/MJD pathogenesis.

Polymorphisms and the relative copy number of the highly mutated parts of mitochondrial genes, including *MT-LT1* (tRNALeu), *MT-TK* (tRNALys), *MT-ND1* (NADH dehydrogenase subunit I), *MT-ATP6/8*(ATP synthase subunit 6/8) and *MT-COII* (Cytochrome C oxidase subunit II), are increasingly being recognized as causing or modifying neurological disease. Thus, our study aims to analyze the polymorphisms and the relative mtDNA copy number variation in mtDNA genes of *MT-LT1*, *MT-ND1*, *MT-CO2*, *MT-TK*, *MT-ATP8* and *MT-ATP6* in 102 SCA3/MJD patients and 100 healthy controls, to determine whether there is a correlation between the mtDNA mutations and clinical phenotypes in SCA3/MJD patients.

## Results

### Polymorphisms analysis of the mtDNA

Among the 102 patients of SCA3/MJD, a total of 24 SNVs were detected in the genes of *MT-LT1*, *MT-ND1*, *MT-CO2*, *MT-TK*, *MT-ATP8* and *MT-ATP6*, two of which were previously reported SNPs, and 22 of which were novel variants (Tab. 1-1). Only 10 SNVs were detected in the 100 healthy controls (Tab. 1-2). The T8772C variant was found in both patient and control groups.

**Tab. 1-1.** SNVs in mtDNA found in SCA3/MJD patient gruop

| Sample no. | Region | Gene | Nucleotide changes | Amino acid changes | Case number | Previously reported |
| --- | --- | --- | --- | --- | --- | --- |
| 1 | 8584 | *MT-ATP6* | G→A | Ala→Thr | 11(10.78%) | no |
| 2 | 8684 | *MT-ATP6* | C→T | Thr→Ile | 2(1.96%) | yes |
| 3 | 8701 | *MT-ATP6* | A→G | Thr→Ala | 48(47.06%) | no |
| 4 | 8860 | *MT-ATP6* | A→G | Thr→Ala | 54(52.94%) | no |
| 5 | 9053 | *MT-ATP6* | G→A | Ser→Asn | 2(1.96%) | no |
| 6 | 8877 | *MT-ATP6* | T→C | null | 3(2.94%) | no |
| 7 | 8793 | *MT-ATP6* | T→C | null | 1(0.98%) | no |
| 8 | 8856 | *MT-ATP6* | G→A | null | 1(0.98%) | no |
| 9 | 8794 | *MT-ATP6* | C→T | His→Tyr | 8(7.84%) | no |
| 10 | 8964 | *MT-ATP6* | C→T | null | 3(2.94%) | no |
| 11 | 8772 | *MT-ATP6* | T→C | null | 1(0.98%) | no |
| 12 | 9123 | *MT-ATP6* | G→A | null | 3(2.94%) | no |
| 13 | 8823 | *MT-ATP6* | T→C | null | 1(0.98%) | no |
| 14 | 8657 | *MT-ATP6* | C→T | Thr→Ile | 1(0.98%) | no |
| 15 | 8718 | *MT-ATP6* | A→G | null | 1(0.98%) | no |
| 16 | 8563 | *MT-ATP6* | A→G | Thr→Ala | 1(0.98%) | no |
| 17 | 8572 | *MT-ATP6* | G→A | Gly→Ser | 1(0.98%) | no |
| 18 | 8974 | *MT-ATP6* | C→T | Leu→Phe | 1(0.98%) | no |
| 19 | 8515 | *MT-ATP8* | C→T | null | 2(1.96%) | no |
| 20 | 8414 | *MT-ATP8* | C→T | Leu→Phe | 2(1.96%) | yes |
| 21 | 8459 | *MT-ATP8* | A→G | Asn→Asp | 3(2.94%) | no |
| 22 | 8440 | *MT-ATP8* | A→G | null | 1(0.98%) | no |
| 23 | 8473 | *MT-ATP8* | T→C | null | 1(0.98%) | no |
| 24 | 8485 | *MT-ATP8* | G→A | null | 1(0.98%) | no |

**Tab. 1-2.** Mutations in mtDNA found in healthy controls

| Sample no. | Region | Gene | Nucleotide changes | Amino acid changes | Case number | Previously reported |
| --- | --- | --- | --- | --- | --- | --- |
| 1 | 3435 | *MT-ND1* | C→T | null | 1(1%) | no |
| 2 | 8772 | *MT-ATP6* | T→C | null | 2(2%) | no |
| 3 | 8945 | *MT-ATP6* | T→C | Met→Thr | 1(1%) | no |
| 4 | 8956 | *MT-ATP6* | A→G | Ile→Val | 1(1%) | no |
| 5 | 8540 | *MT-ATP8* | T→C | Leu→Pro | 1(1%) | no |
| 6 | 8730 | *MT-ATP6* | A→G | null | 3(3%) | no |
| 7 | 8785 | *MT-ATP6* | C→T | Leu→Phe | 1(1%) | no |
| 8 | 8940 | *MT-ATP6* | C→T | null | 2(2%) | no |
| 9 | 8970 | *MT-ATP6* | C→T | null | 1 | no |
| 10 | 9080 | *MT-ATP6* | A→G | Asn→Ser | 1 | no |

### Analysis of the relative mtDNA copy number

There was no difference in the mtDNA copy number (SCA3/MJD: 93.20, healthy controls: 89.66, P>0.05, Fig.1-1) between SCA3/MJD patients and healthy controls. In the SCA3/MJD patient group, the relative mtDNA copy number had a negative correlation with the number of CAG repeats (r=-0.210, P < 0.05, Fig.1-2) but did not correlate with the age at testing, the age of onset, disease duration, ICARS or SARA score (Tab.2-2).

**Fig. 1-1.**
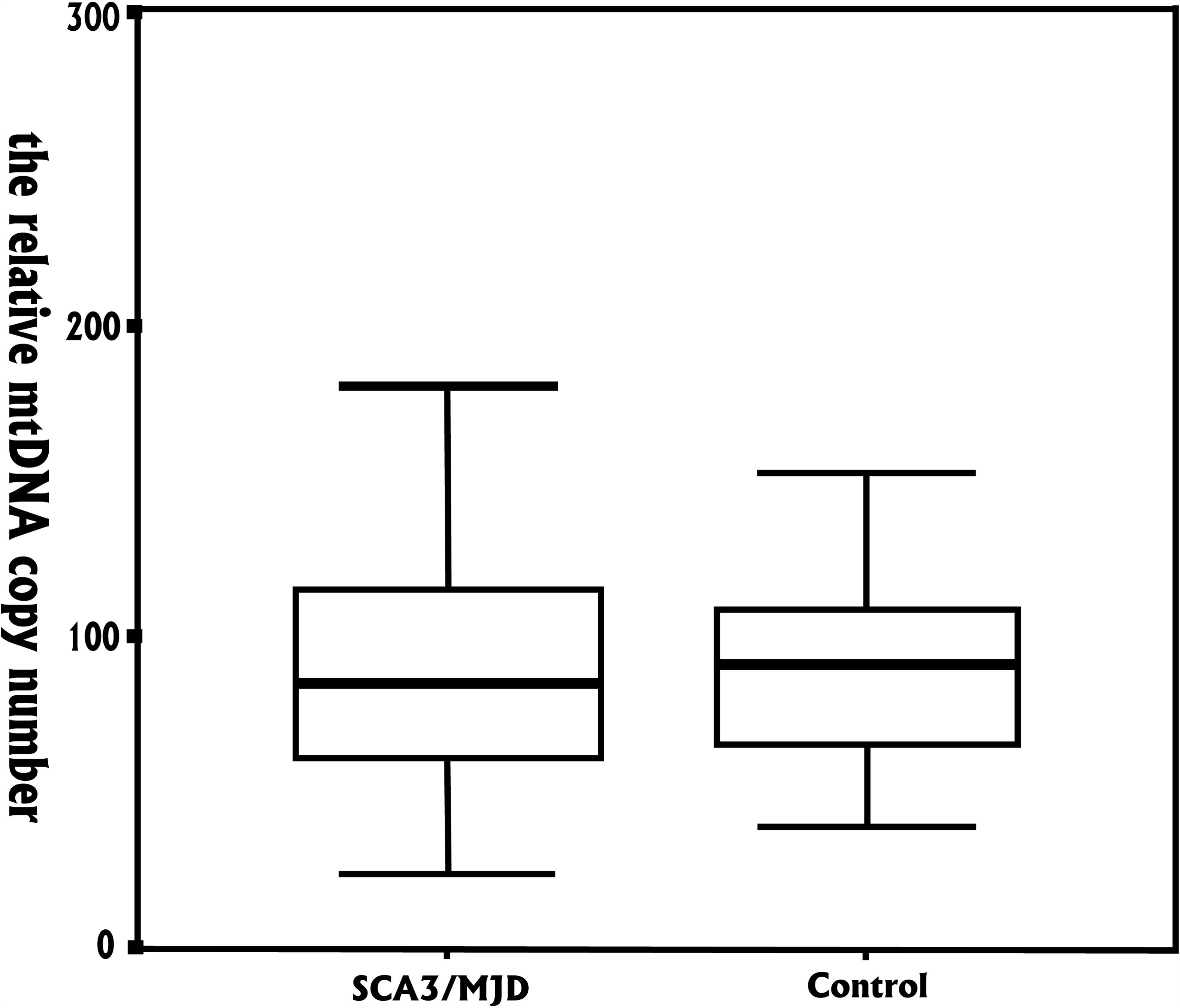
Relative mtDNA copy number in SCA3/MJD and controls.

**Fig. 1-2.**
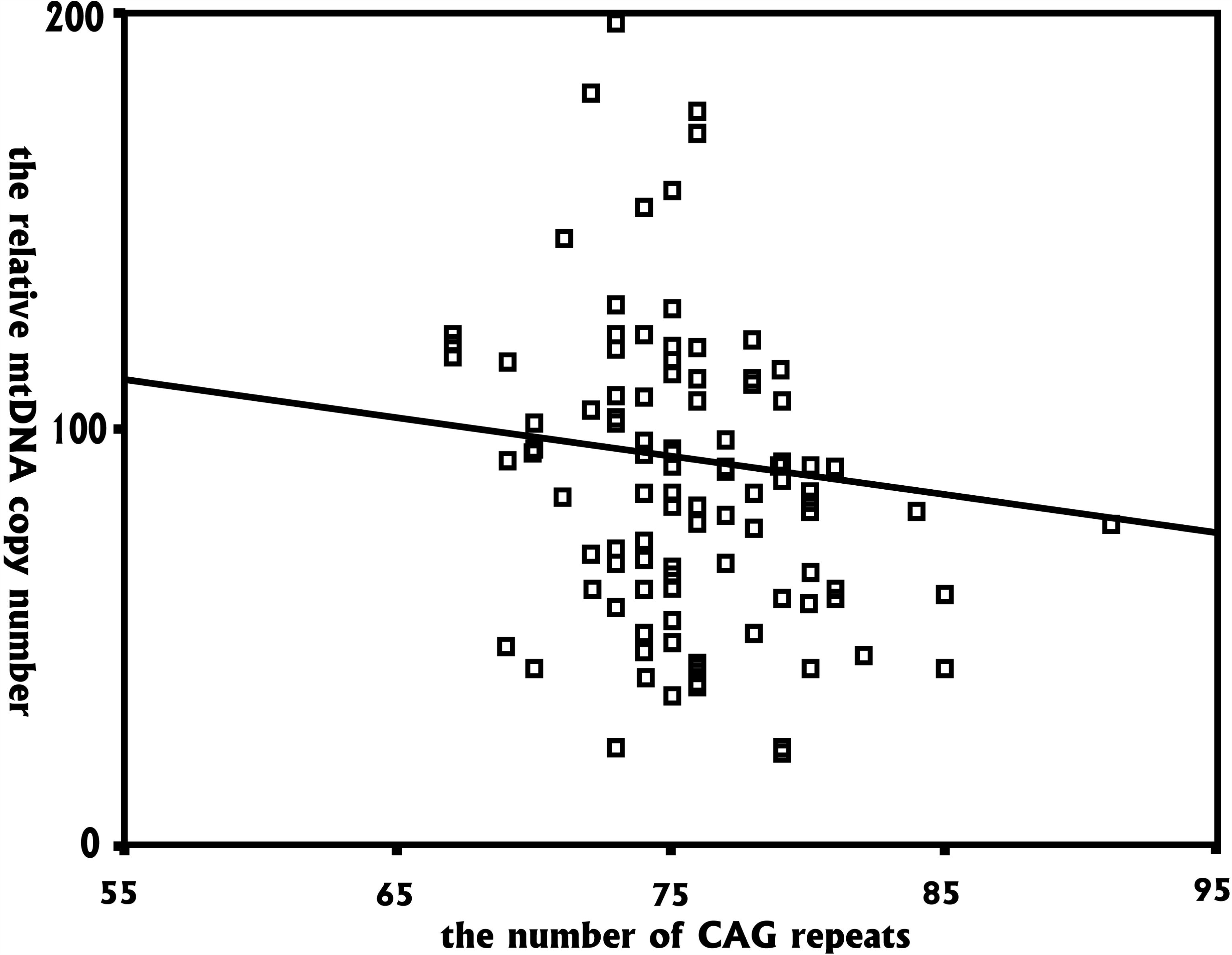
Negative correlation of the relative mtDNA copy number with the number of.

**Tab. 2-1.**
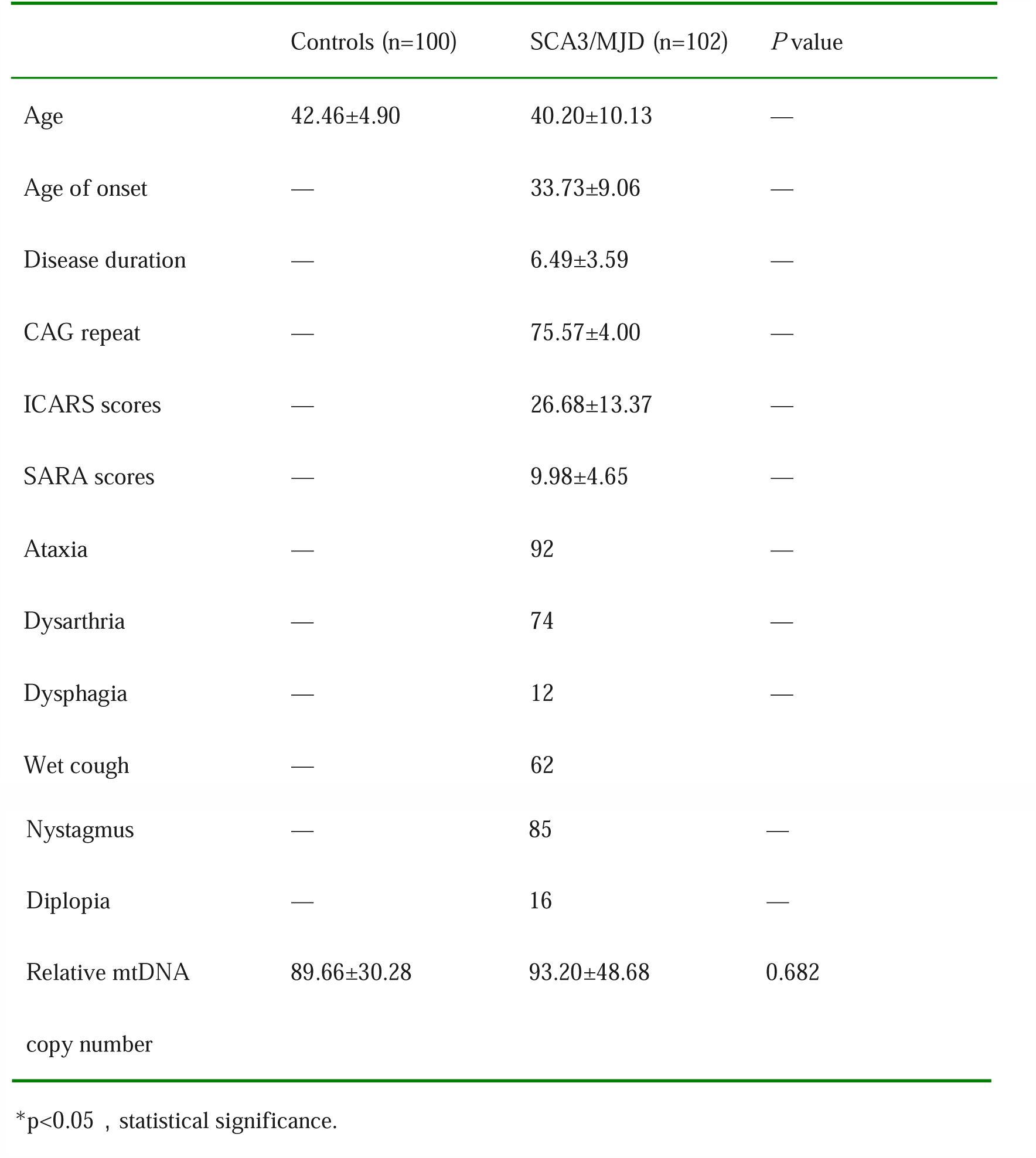
Clinical and molecular parameters of SCA3/MJD patients and controls

|  | Controls (n=100) | SCA3/MJD (n=102) | *P* value |
| --- | --- | --- | --- |
| Age | 42.46±4.90 | 40.20±10.13 | — |
| Age of onset | — | 33.73±9.06 | — |
| Disease duration | — | 6.49±3.59 | — |
| CAG repeat | — | 75.57±4.00 | — |
| ICARS scores | — | 26.68±13.37 | — |
| SARA scores | — | 9.98±4.65 | — |
| Ataxia | — | 92 | — |
| Dysarthria | — | 74 | — |
| Dysphagia | — | 12 | — |
| Wet cough | — | 62 | — |
| Nystagmus | — | 85 | — |
| Diplopia | — | 16 | — |
| Relative mtDNA copy number | 89.66±30.28 | 93.20±48.68 | 0.682 |

\* $p < 0.05$ , statistical significance.

**Tab. 2-2.**
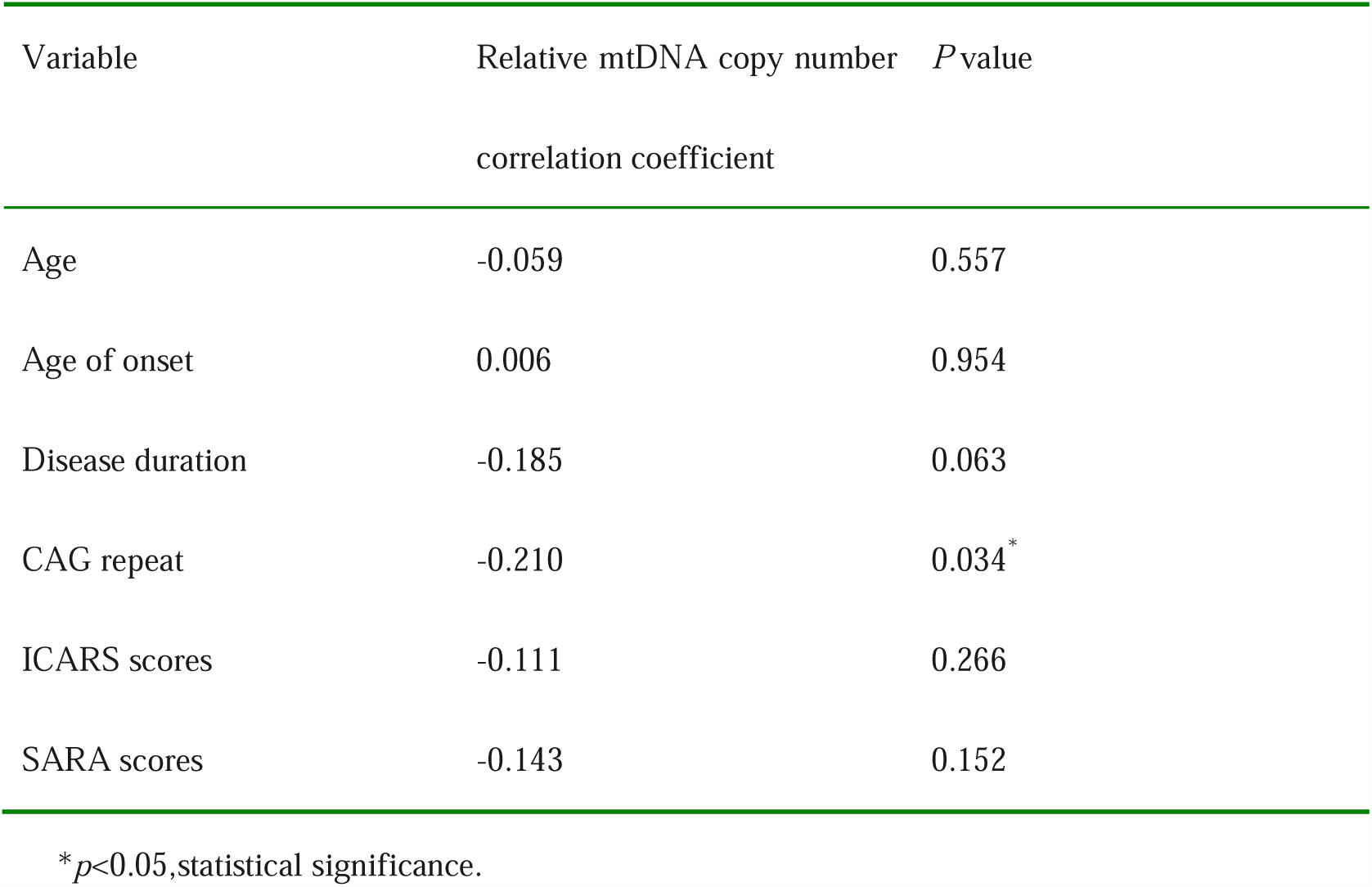
Spearman test of relative mtDNA copy number with various parameters in SCA3/MJD patients

## Discussion

Mitochondrial DNA is considered as the smallest chromosome, containing only approximately 16,569 base pairs and coding for 37 genes. It is located in the mitochondria, which are completely separated from the nuclear genome. Mutations in mtDNA can cause failure of oxidative phosphorylation and the ATP synthesis, contributing to the oxidative stress and neuronal death found in many neurodegenerative conditions(Zeviani *et al.* 2012)^9^[12], such as Alzheimer’s disease(Sheng *et al.* 2012), Parkinson’s disease(Bose and Beal 2016)^11^[14], Huntington’s disease and diabetes(Trushina and Mcmurray 2007; Schapira 2008). The pathogenesis of SCA3/MJD is complex, and the role of oxidative stress, if any, is unclear. Our research aimed to discover mtDNA mutations that might explain some of the phenotypic variability in SCA3/MJD.

Many groups have reported mtDNA mutations in various hereditary ataxia syndromes. Morgan-Hughes(Pulkes *et al.* 1999) and his partners reported 4 patients had ataxia symptom in 14 MELAS (mitochondrial encephalomyopathy with lactic acidosis and stroke-like episodes) patients with mtDNA A3243G mutation. Also, MERRF (myoclonus epilepsy ragged-red fibers) patients with A8344G mutation of mtDNA were reported to be ataxic. Neuropathy-ataxia-retinitis pigmentosa syndrome patients also have T8993C and T8993G mtDNA mutations.

Mitochondrial morphological change is also considered as an indicator of mitochondrial dysfunction. Electron microscopy of liver tissue revealed mitochondrial morphological changes in a patient with SCA7 (Han *et al.* 2010).In our study, we analyzed the mutations in several commonly reported mtDNA genes, including *MT-LT1*, *MT-ND1*, *MT-CO2*, *MT-TK*, *MT-ATP8* and *MT-ATP6*. Interestingly, we found 22 new SNVs in ATP8 and ATP6. Among these SNVs, the variant rates of A8701G and A8860G were high (47.06% and 52.94%, respectively), and both cause replacement of the threonine with alanine. Threonine is neutral polar amino acid, and alanine is neutral non-polar amino acid. Thus, the threonine to alanine mutation may cause a structural change in the translated product, but whether it will cause a failure in energy metabolism is still unclear. Only when the mutation rate of the mtDNA reaches a certain threshold do the cells and organelles may show functional changes(Chen *et al.* 2015). In our study, the SNV rate in mtDNA is much higher in SCA3/MJD patients than the normal controls, which may indicate a role for mtDNA mutation in SCA3 pathogenesis. It is unclear whether these mutations are modifiers of disease, or whether they are a consequence of changes caused by repeat expansion in the ataxin-3 gene.

In previous studies, it was shown that mtDNA copy numbers from peripheral leucocytes in MELAS patients or MERRF patients correlated negatively with the mtDNA mutation rate(Chabi *et al.* 2003). In an SCA3/MJD cell model, decreased antioxidant activity lead to increased mtDNA damage, including a decreased mtDNA copy number and the loss of 4977 base pairs2. In another cohort of SCA and Huntington’s disease patients, Bing-Wen Soong showed that the decrease in mtDNA copy number from peripheral white blood cells negatively correlates with the number of CAG repeats, and proposed that mtDNA copy number could be a marker for these trinucleotide repeat diseases(Liu *et al.* 2008). We analyzed the mtDNA copy numbers variant from peripheral white blood cells in both SCA3/MJD patients and healthy controls, and found a negative correlation between the relative mtDNA copy number and the number of CAG repeats in SCA3/MJD patients. There was no correlation with the age at testing, the age of onset, disease duration, ICARS scores or SARA scores, and we did not observe a significant difference in relative mtDNA copy number between SCA3/MJD patients and healthy controls.

In summary, our study demonstrated that the frequency of mutated mtDNA in SCA3/MJD patients was higher than in the healthy group, suggesting that poly-Q repeat length in this disease may be associated with mutations in mtDNA. The mtDNA relative copy number in SCA3/MJD patients was not significantly different compared with the healthy control group, indicating that mtDNA copy number is not a biomarker for disease severity in SCA3/MJD patients.

## Subjects and methods

### Patients and controls

One hundred and two unrelated SCA3/MJD patients (53 males and 49 females), confirmed molecularly, and 100 unrelated healthy controls (50 males and 50 females), matched by age and sex, were recruited for the study from Xiangya Hospital, Central South University, China between June 2007 and June 2015. The study was approved by the Expert Committee of Xiangya Hospital of the Central South University in China (equivalent to an Institutional Review Board). Written informed consent was obtained from all subjects. None of the subjects had a history of exposure to occupational carcinogens, familial history of cancer, benign or malignant tumors, smoking, drinking or other unhealthy habits, hypertension, cardiovascular diseases, inflammatory disease, kidney disease or diabetes other than SCA3/MJD. Informed consent was obtained from all participants. A 5-10 ml sample of blood was collected in heparin after venipuncture. Total DNA, extracted with phenol–chloroform, was stored at −20 centigrade.

### Polymorphisms analysis of the mtDNA

The two pairs of primers (Tab. 3) used to amplify and sequence the mtDNA SNVs were previously published(Safaei *et al.* 2009). The first pair is ONP25 and ONP185, used for amplifying *MT-TK*, *MT-ATP8*, *MT-ATP6* and a portion of the *MT-CO2* gene in the mtDNA (approximately 1078 bp). The another pair is ONP82 and ONP164, used for amplifying *MT-LT1* and a portion of the *MT-ND1* gene in the mtDNA (about 363 bp). The PCR conditions were as follows: predenaturation at 94°C for 5 minutes, 35 cycles of 94°C for 1 minute, 60°C for 1 minute, 72°C for 1 minute (363 bp fragment for 35 seconds), followed by a final extension at 72°C for 10 minutes. The amplified fragments were purified and sequenced by Sanger sequencing on a 3730 Genetic Analyzer sequencing machine (Applied Biosystems). SNPs that have been previously reported were identified by comparison with the Polymorphism Database in NCBI and TSC (bSNP Http://www.ncbi.nlm.nih.gov/SNP/; TSC http://snp.cshl.org/), and mutations that have been previously reported were identified by comparison with the Human Gene Mutation Database (http://www.hgmd.cf.ac.uk/ac/index.php).

**Tab.3.**
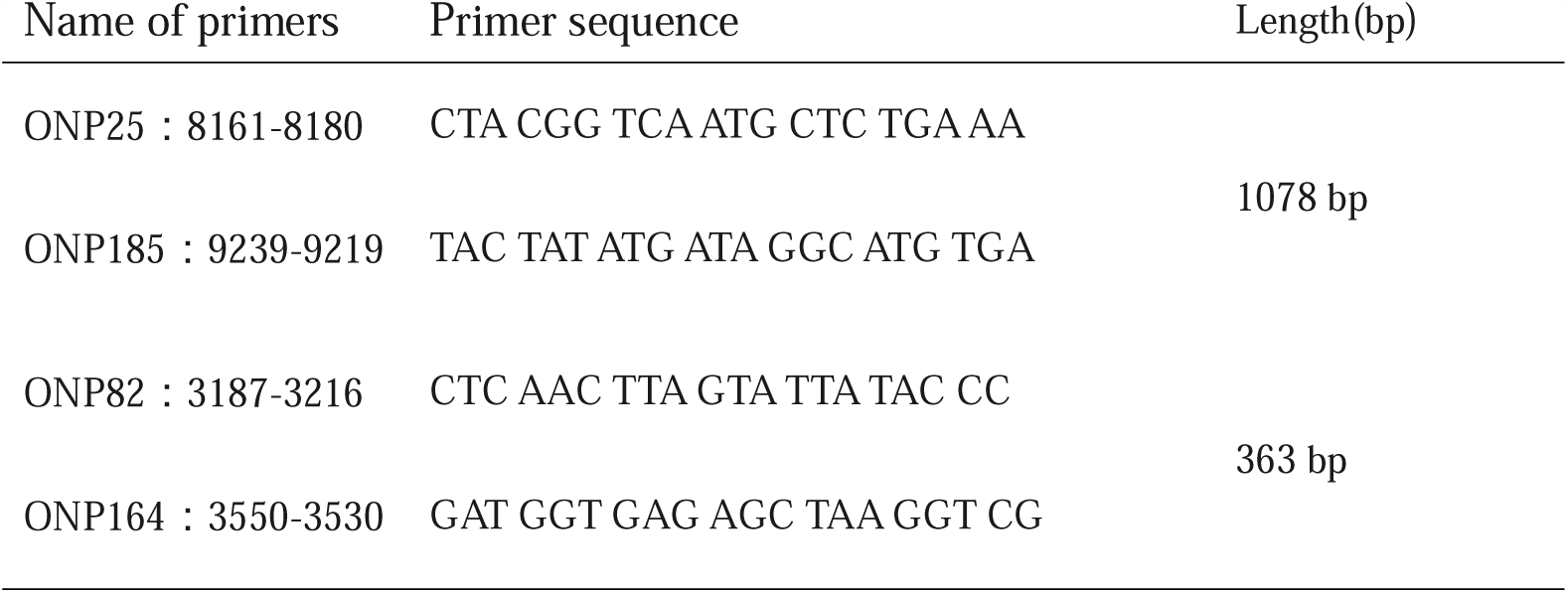
Primers used for screening the mutation of the mtDNA

### Analysis of the relative mtDNA copy number

The relative mtDNA copy number was measured using real-time PCR for the *ND1* gene and normalized against the reference gene *β-globin*. Pooled DNA from the Health Study participants served as the reference DNA pool, used to create a fresh standard curve on which the system was calibrated, which ranged from 0.16 to 20 ng/μL. All samples were assessed in triplicate, and the average of all three measurements was calculated. We used primers from Liu et al(Liu *et al.* 2003) for this study(Tab.4). The PCR conditions for *β-globin* and *ND1* were 95°C for 30 seconds, 35 cycles of 95°C for 5 seconds, 57°C for 20 seconds and 72°C for 15 seconds.

**Tab.4.**
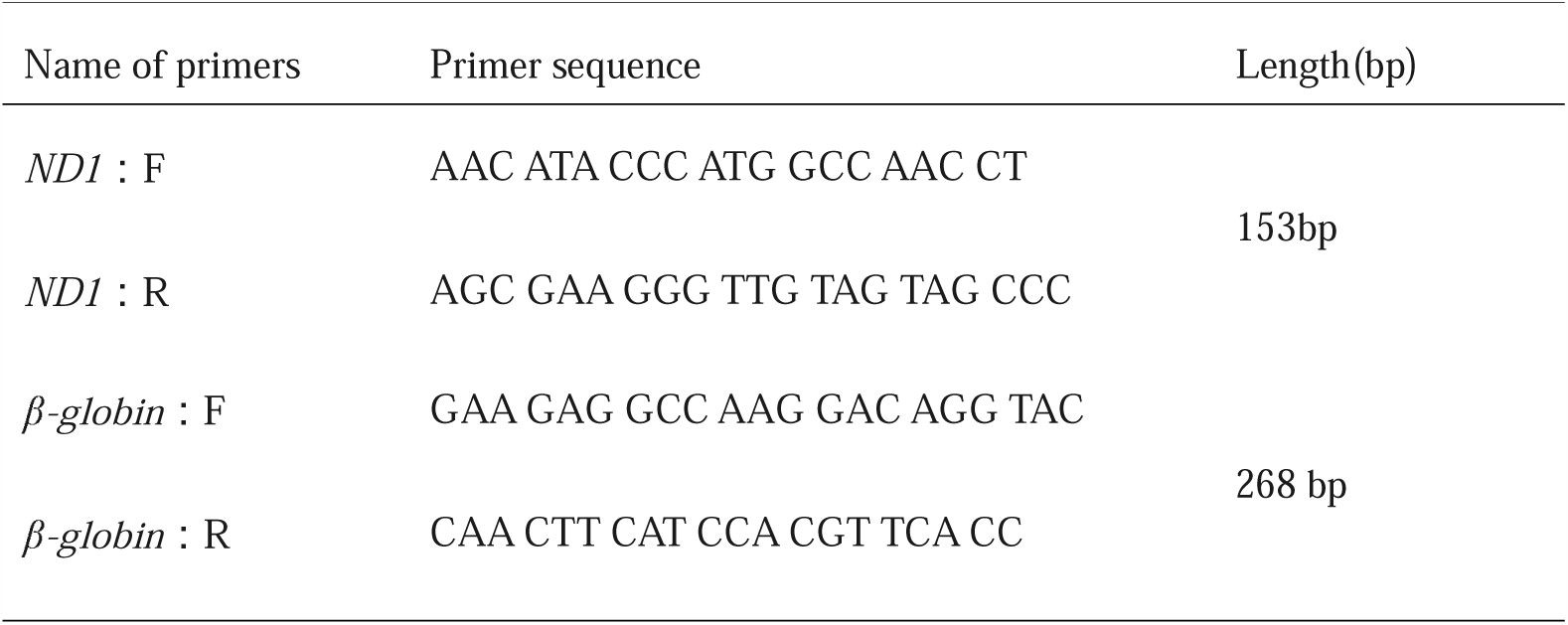
Primers used for real-time PCR analysis

### Statistical analysis

Statistical analysis was carried out with SPSS 22.0 and Excel 2010. All data were expressed as the means ± standard deviation. The Mann-Whitney U test was used for comparison of the relative mtDNA copy number between SCA3/MJD patients and healthy controls, and correlations of relative mtDNA copy number with CAG repeat, age at detection, age of onset, disease duration, ICARS scores and SARA scores were assessed by the Spearman correlation test. The data were considered significant when the P value was less than 0.05.

## Acknowledgments

We are grateful to the participating patients for their involvement. This study was supported by the Program of National Natural Science Foundation of China (#81671120, 81300981, 81400938, 81250015), National Department Public Benefit Research Foundation (#201302001), The innovation project of Central South University graduate students (#502210004). Innovation driven program of Central South University (#506010108).

We thank Fay Alexander thank for the language help on writing and editing of the manuscript.

## Author Contributions

Zhen Liu performed research, analyzed the data, and wrote the paper. Jie Zhou performed research and analyzed the data. Xiaomeng Yin modify the article. Shuyin Shi and Weining Sun performed research. Hong jiang, Lu Shen and Beisha Tang supervised the study. Junling Wang directed the overall research and wrote the manuscript.

## References

Bose, A., and M. F. Beal, 2016 Mitochondrial dysfunction in Parkinson’s disease. J Neurochem.

Casali, C., G. M. Fabrizi, F. M. Santorelli, G. Colazza, M. Villanova et al., 1999 Mitochondrial G8363A mutation presenting as cerebellar ataxia and lipomas in an Italian family. Neurology 52: 1103–1104.

Chabi, B., B. Mousson de Camaret, H. Duborjal, J. P. Issartel and G. Stepien, 2003 Quantification of mitochondrial DNA deletion, depletion, and overreplication: application to diagnosis. Clin Chem 49: 1309–1317.

Chen, Y., W. D. Parker, H. Chen and K. Yang, 2015 Aberrant mitochondrial RNA in the role of aging and aging associated diseases. Med Hypotheses 85:178–182.

Chinnery, P. F., D. T. Brown, K. Archibald, A. Curtis and D. M. Turnbull, 2002 Spinocerebellar ataxia and the A3243G and A8344G mtDNA mutations. J Med Genet 39: E22.

Han, Y., B. Deng, M. Liu, J. Jiang, S. Wu et al., 2010 Clinical and genetic study of a Chinese family with spinocerebellar ataxia type 7. Neurol India 58: 622–626.

Lee, Y. C., Y. C. Lu, M. H. Chang and B. W. Soong, 2007 Common mitochondrial DNA and POLG1 mutations are rare in the Chinese patients with adult-onset ataxia on Taiwan. J Neurol Sci 254: 65–68.

Liu, C. S., W. L. Cheng, S. J. Kuo, J. Y. Li, B. W. Soong et al., 2008 Depletion of mitochondrial DNA in leukocytes of patients with poly-Q diseases. J Neurol Sci 264: 18–21.

Liu, C. S., C. S. Tsai, C. L. Kuo, H. W. Chen, C. K. Lii et al., 2003 Oxidative stress-related alteration of the copy number of mitochondrial DNA in human leukocytes. Free Radic Res 37:1307–1317.

Maruyama, H., S. Nakamura, Z. Matsuyama, T. Sakai, M. Doyu et al., 1995 Molecular features of the CAG repeats and clinical manifestation of Machado-Joseph disease. Hum Mol Genet 4: 807–812.

Nobrega, C., I. Nascimento-Ferreira, I. Onofre, D. Albuquerque, H. Hirai et al., 2013 Silencing mutant ataxin-3 rescues motor deficits and neuropathology in Machado-Joseph disease transgenic mice. PLoS One 8: e52396.

Padiath, Q. S., A. K. Srivastava, S. Roy, S. Jain and S. K. Brahmachari, 2005 Identification of a novel 45 repeat unstable allele associated with a disease phenotype at the MJD1/SCA3 locus. Am J Med Genet B Neuropsychiatr Genet 133b: 124–126.

Pareyson, D., P. Saveri and G. Piscosquito, 2014 Charcot-Marie-Tooth Disease and Related Hereditary Neuropathies: From Gene Function to Associated Phenotypes. Curr Mol Med.

Petrozzi, L., G. Ricci, N. J. Giglioli, G. Siciliano and M. Mancuso, 2007 Mitochondria and neurodegeneration. Biosci Rep 27: 87–104.

Pulkes, T., L. Eunson, V. Patterson, A. Siddiqui, N. W. Wood et al., 1999 The mitochondrial DNA G13513A transition in ND5 is associated with a LHON/MELAS overlap syndrome and may be a frequent cause of MELAS. Ann Neurol 46: 916–919.

Ramos, A., N. Kazachkova, F. Silva, P. Maciel, A. Silva-Fernandes et al., 2015 Differential mtDNA damage patterns in a transgenic mouse model of Machado-Joseph disease (MJD/SCA3). J Mol Neurosci 55: 449–453.

Safaei, S., M. Houshmand, M. M. Banoei, M. S. Panahi, S. Nafisi et al., 2009 Mitochondrial tRNALeu/Lys and ATPase 6/8 gene variations in spinocerebellar ataxias. Neurodegener Dis 6:16–22.

Schapira, A. H., 2008 Mitochondrial dysfunction in neurodegenerative diseases. Neurochem Res 33: 2502–2509.

Sheng, B., X. Wang, B. Su, H. G. Lee, G. Casadesus et al., 2012 Impaired mitochondrial biogenesis contributes to mitochondrial dysfunction in Alzheimer’s disease. J Neurochem 120: 419–429.

Trushina, E., and C. T. McMurray, 2007 Oxidative stress and mitochondrial dysfunction in neurodegenerative diseases. Neuroscience 145:1233–1248.

Waldbaum, S., and M. Patel, 2010 Mitochondria, oxidative stress, and temporal lobe epilepsy. Epilepsy Res 88: 23–45.

Yu, Y. C., C. L. Kuo, W. L. Cheng, C. S. Liu and M. Hsieh, 2009 Decreased antioxidant enzyme activity and increased mitochondrial DNA damage in cellular models of Machado-Joseph disease. J Neurosci Res 87: 1884–1891.

Zeviani, M., A. Simonati and L. A. Bindoff, 2012 Ataxia in mitochondrial disorders. Handb Clin Neurol 103: 359–372.

